# GALA: gap-free chromosome-scale assembly with long reads

**DOI:** 10.1101/2020.05.15.097428

**Authors:** Mohamed Awad, Xiangchao Gan

## Abstract

High-quality genome assembly has wide applications in genetics and medical studies. However, it is still very challenging to achieve gap-free chromosome-scale assemblies using current workflows for long-read platforms. Here we propose GALA (**Ga**p-free **l**ong-read **a**ssembler), a chromosome-by-chromosome assembly method implemented through a multi-layer computer graph that identifies mis-assemblies within preliminary assemblies or chimeric raw reads and partitions the data into chromosome-scale linkage groups. The subsequent independent assembly of each linkage group generates a gap-free assembly free from the mis-assembly errors which usually hamper existing workflows. This flexible framework also allows us to integrate data from various technologies, such as Hi-C, genetic maps, a reference genome and even motif analyses, to generate gap-free chromosome-scale assemblies. We *de novo* assembled the *C. elegans* and *A. thaliana* genomes using combined Pacbio and Nanopore sequencing data from publicly available datasets. We also demonstrated the new method’s applicability with a gap-free assembly of a human genome with the help a reference genome. In addition, GALA showed promising performance for Pacbio high-fidelity long reads. Thus, our method enables straightforward assembly of genomes with multiple data sources and overcomes barriers that at present restrict the application of *de novo* genome assembly technology.

## Introduction

*De novo* genome assembly has wide applications in plant, animal and human genetics. However, it is still very challenging for long-read platforms, such as Nanopore and Pacbio, to provide chromosome-scale sequences [1, 2]. To date, numerous *de novo* assembly tools have been developed to obtain longer and more accurate representative sequences from raw sequencing data [3–5]. In most studies, however, assemblies by these tools comprise hundreds or even thousands of contigs. To produce chromosome-scale assembly, various technologies, such as Hi-C, genetic maps or even a reference genome, have been increasingly used to anchor contigs into big scaffolds[6, 7]. As a consequence, the final genome assembly usually contains numerous gaps, and sometimes, are also plagued with mis-assemblies [8, 9].

Gaps and mis-assemblies in a genome assembly can seriously undermine genomic studies. For example, a lot of sequence alignment tools have much lower performances when query sequences contain gaps [10, 11]. In intraspecific genome comparisons, large gaps not only significantly increase the possibility of failure to detect long structure variants, but also produce inaccurate results of gene annotation [12, 13]. Moreover, gaps and mis-assemblies have been reported to accounts for a large number of gene model errors in existing genome assembly studies [14, 15].

In this study, we report on GALA (**Ga**p-free **l**ong-read **a**ssembler), a scalable method, for gap-free chromosome-scale assembly (**Fig. 1**). GALA separates two steps: firstly, it identifies multiple linkage groups in the genome, each representing a single chromosome, and it describes chromosome structure with raw reads and assembled contigs from multiple *de novo* assembly tools; secondly, it assembles each linkage group by integrating results from multiple assembly tools and inference from raw reads. Moreover, our method can also exploit the information derived from Hi-C data to obtain chromosome-scale linkage groups in studies even with a complicated genome structure or those with low sequencing quality. Of note is that our method can be easily extended to incorporate other sources of information, such as genetic maps or even genome synteny information. We demonstrate the utility of GALA by gap-free and chromosome-scale assemblies of Pacbio or Nanopore sequencing data from two publicly available datasets for which the original assembly contains large gaps and a number of unanchored scaffolds. Notably, our approach significantly outperforms existing algorithms in both datasets. Further, we also demonstrate the application of our method to assemble a human genome with the help a reference genome using Pacbio HiFi sequencing data.

**Figure 1.**
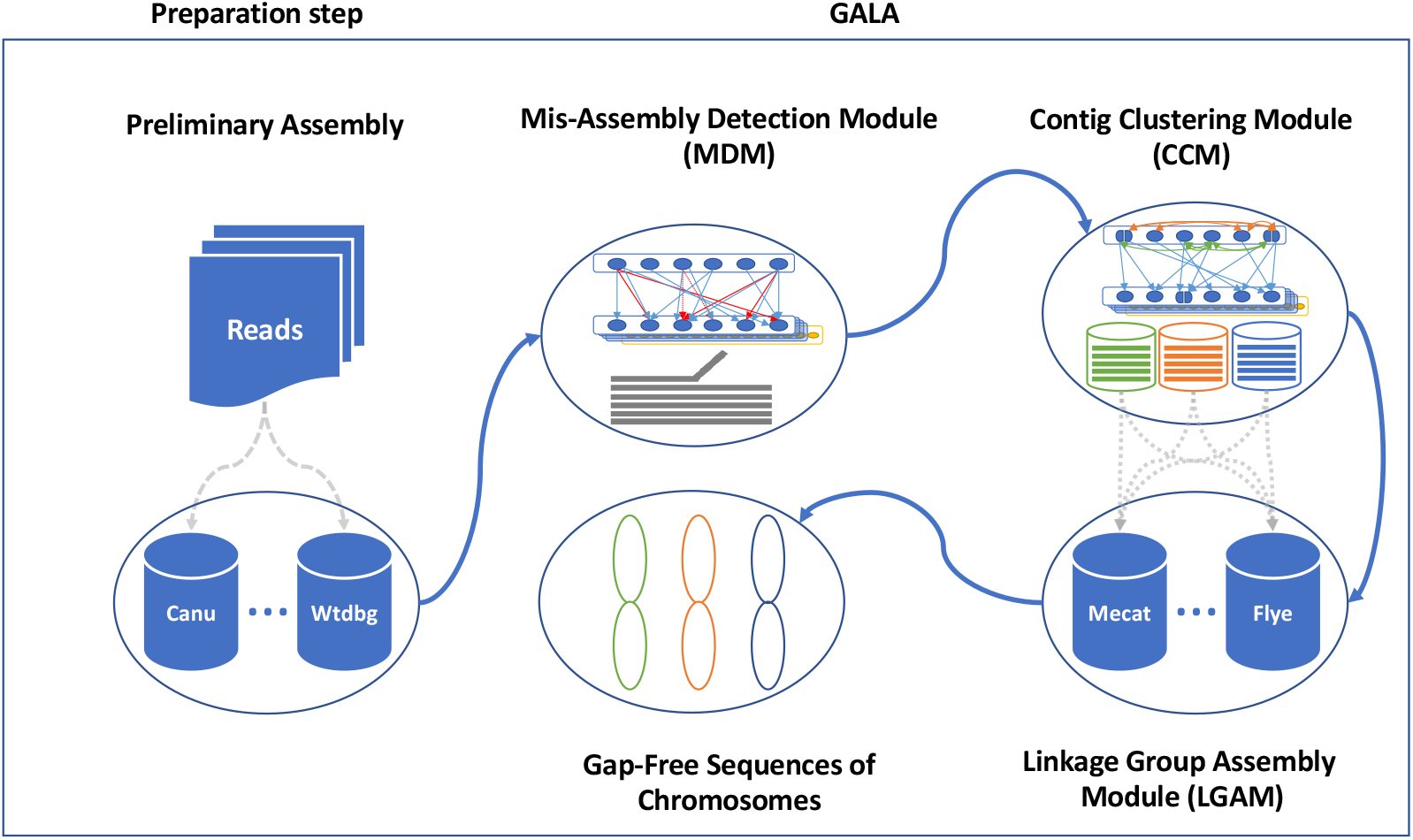
Overview of GALA. After *de novo* assembling with various tools, preliminary assemblies and raw reads are encoded into a multi-layer computer graph. Mis-assemblies are identified with MDM by browsing through the inter-layer information. The split nodes are clustered into multiple linage groups by the CCM. Each linkage group is assembled independently using LGAM to achieve the final gap-free sequences of chromosomes.

## Results

### Overview of the GALA framework

GALA exploits the information from multiple *de novo* assembly tools and raw reads, as well as other information sources, such as Hi-C, genetic maps or even a reference genome, if they exist. In GALA, a batch of *de novo* assembly tools are selected to first create preliminary assemblies. These preliminary assemblies and raw reads are then aligned against each other. We use a multi-layer computer graph to model the GALA, with each assembly encoded as one layer, together with an extra layer representing the raw reads. Inside each layer, a contig (or a read in the raw-read layer) is encoded as a graph node. GALA browses through the reciprocal alignments and creates two types of edges. Any contradictory information between multiple assemblies or raw reads is recorded as a cross-layer edge. Inside each layer, if two nodes both partially overlap with the same node inside a different layer, a within-layer edge is created between them (**Fig. 2**).

**Figure 2.**
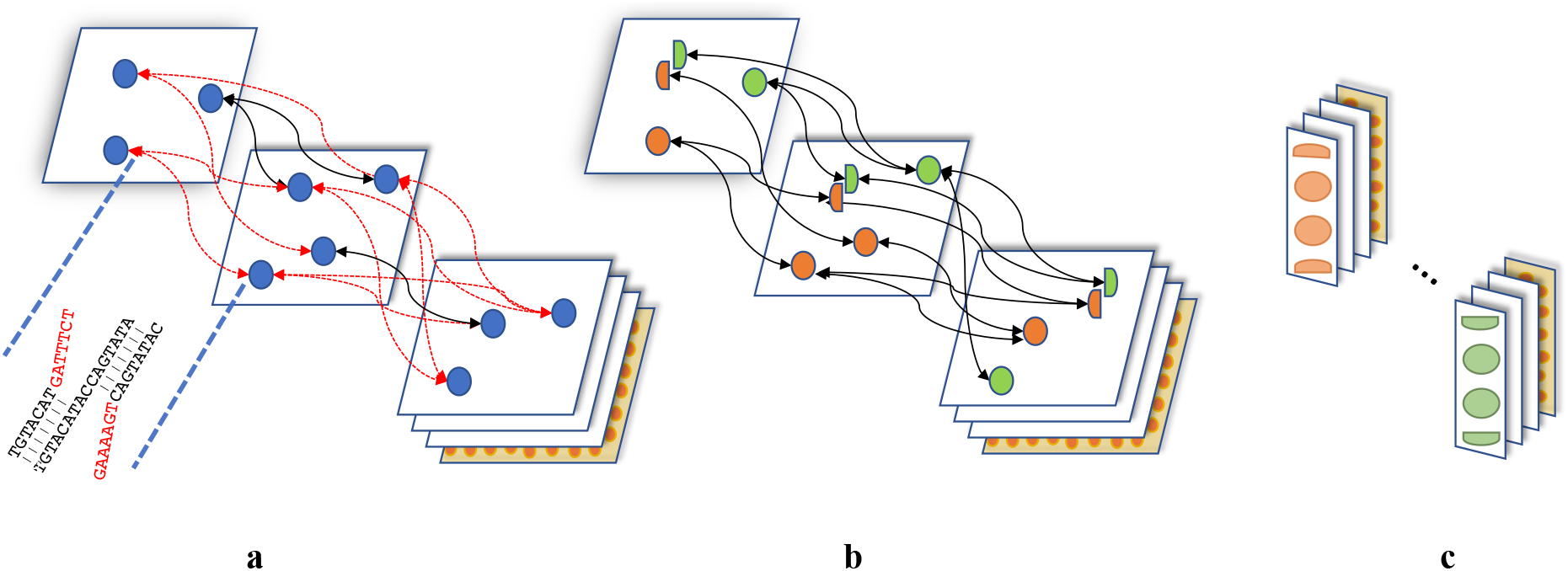
Illustration of a multi-layer computer graph in GALA. (a) The preliminary assemblies and raw reads are aligned against each other and encoded into a multi-layer graph. Conflicted alignments are indicated in red. (b) The conflicted alignments are removed iteratively by splitting the nodes involved and new edges are assigned accordingly. (c) Nodes are clustered into different linkage groups.

Depending on the sequencing quality and complexity of the genome structure, existing assembly tools usually exhibit different performances in terms of the number of misassembled contigs and N50. To prevent the spread of errors, we developed a mis-assembly detection module (MDM), which works by cutting out the contradictory cross-layer edges by splitting nodes (Methods). After removing contradictory cross-layer links, the contig-clustering module (CCM) pools the linked nodes within different layers and those inside the same layer into different linkage groups, usually each representing a chromosome (Methods). In several experiments, we identified orphan contigs. Interestingly, most of them come from external sources such as bacterial or sample contamination.

The successful partitioning of existing preliminary assemblies and raw reads into separate linkage groups allows us to essentially perform a chromosome-by-chromosome assembly. The raw reads from each linkage group are extracted and assembled with multiple assembly tools and merged together if necessary. For those tools which take corrected reads as input, we correct reads using suggested methods. Interestingly, we found that chromosome-by-chromosome assembly always provides better performance, especially for the repetitive fragments, while the improvement of read-correction with chromosome-by-chromosome analysis is negligible. GALA also provides a fast mode, where the consensus assembly for each chromosome is obtained by merging the assembled contigs within the linkage group without working on raw reads. However, in many cases, the fast mode generated gapped assemblies, thereby highlighting the distinct advantage of the chromosome-by-chromosome assembly strategy over existing tools.

### *Caenorhabditis elegans* genome assembly

We used a publicly available dataset for *Caenorhabditis elegans* VC2010. The dataset was generated on the Pacbio platform with a 290X coverage along with an extra 32X coverage of Nanopore sequence [16]. As no current assembly tools support pooled sequencing data from Pacbio and nanopore platforms, we used both datasets separately to generate preliminary assemblies (**Supplementary Fig. 1**). Preliminary assemblies were generated using Canu, Flye, Mecat2/Necat, Miniasm and Wtdbg2 (Methods). Among all our preliminary assemblies, the one produced by Pacbio-Flye showed the smallest number of contigs, with 41 contigs for 102Mbp of overall sequence.

We applied GALA to the raw reads and the preliminary assemblies. The numbers of mis-assemblies in each preliminary assembly derived by the MDM algorithm ranged from 0 to 19. After removing the mis-scaffolds through the node-splitting operation, GALA modelled the input into 14 independent linkage groups. Seven of them are from short continuous contigs and the others represent individual chromosomes or chromosome arms. After removing the four contigs from bacterial contamination or organelle DNA, we were able to merge the graph further into six linkage groups through telomeric motif analyses (**Supplementary Fig. 2** and Methods). Of note is that the integrative assembly of each linkage group generated gap-free complete sequences for all six chromosomes.

We performed additional analyses to test the performance of our assembly using the Hi-C dataset generated by the same group. By aligning the Hi-C data against our assembly using BWA-MEM, then detecting the potential mis-assemblies using Salsa [17], no mis-assemblies were revealed. Salsa also supported the merging of two linkage groups suggested by the telomeric motif analyses in our assembly. For comparison, we also applied Salsa with Hi-C data to the best preliminary assembly from Flye with Pacbio data. This Flye/Hi-C assembly contains eight scaffolds and 14 unanchored contigs after excluding those from sample contamination. We observed 13 spanned gaps in the Flye/Hi-C assembly with the two largest gaps being 4.95 Mbp and 1.59 Mbp (**Fig 3**).

**Figure 3.**
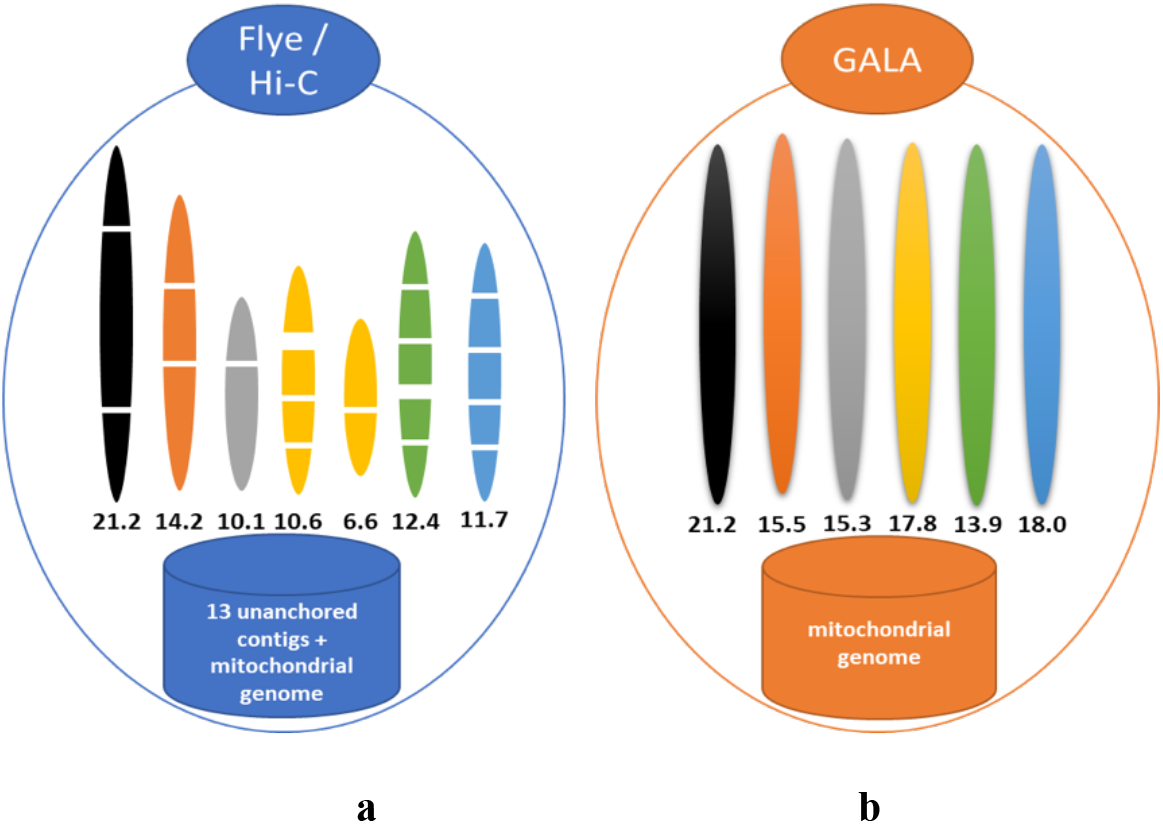
Comparison of Flye assembly with Hi-C scaffolding and GALA assembly of long reads of the *C. elegans* genome. (a) The Flye assembly with Hi-C scaffolding contains numerous gaps and 13 unanchored contigs in the assembly. (b) GALA produces gap-free assembly for each chromosome. Note this is not a fair comparison since GALA did not use Hi-C data in this assembly.

We polished our assembly using Pacbio and Illumina short reads and then compared it to the published assembly. The evaluation from Busco 3.0.0 indicated that our assembly successfully assembled two more genes. The alignment of Illumina short reads against our assembly also reveals a better alignment rate as well as fewer variants (**Table 1** and **Supplementary Fig. 3**).

**Table 1.** The assembly performance evaluation of GALA with Illumina short reads.

|  | REF(N2) | VC2010 | GALA assembly |
| --- | --- | --- | --- |
| **Mapped reads** | 130,604,410 | 130,639,345 | 130,652,108 |
| **Unmapped reads** | 4,568,540 | 4,533,605 | 4,520,842 |
| **Variants** | 17,385 | 14,839 | 14,169 |
| **SNPs** | 16,179 | 14,167 | 13,701 |
| **Deletions** | 412 | 282 | 124 |
| **Insertions** | 794 | 390 | 344 |
| **Indels** | 1,206 | 672 | 468 |

### Arabidopsis *thaliana* genome assembly

We assembled *Arabidopsis thaliana* accession KBS-Mac-74 by combinatory analysis of two publicly available datasets using GALA, one is from Pacbio with a 58X coverage and the other is from Nanopore with a 28X coverage [18]. We used both datasets separately to generate preliminary assemblies. Both raw reads and corrected reads by Canu and Mecat2/Necat (**Supplementary Fig. 4**) were used as input for Canu, Flye, Mecat2/Necat, Miniasm and Wtdbg2 assemblers (Methods).

**Figure 4.**
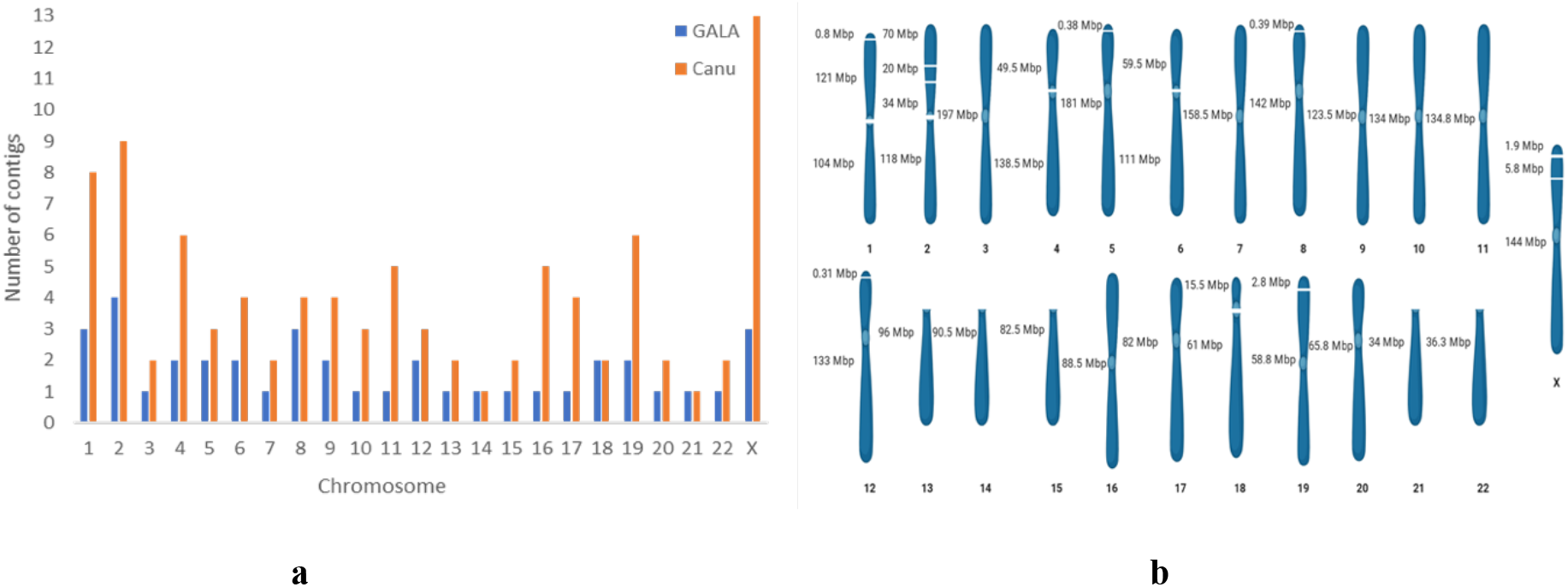
The performances of the human genome assembly GALA. (a) Comparison of the number of contigs in assemblies by canu and GALA. (b) A cartoon presentation of each chromosome assembled by GALA, with the lengths of contigs labelled.

GALA analyses on the raw reads and the preliminary assemblies highlighted a number of mis-assemblies for each preliminary assembly, ranging from 1-18. GALA modelled the input into 14 independent linkage groups. Among them, one from the mitochondrial genome, one from the chloroplast genome and three are continuous fragments of 1.6 Kbp, 7.6 Kbp and 18.5 Kbp. The remaining ten linkage groups represent a chromosome arm each. Previous studies have indicated that the Pacbio and Nanopore platforms seldomly sequence through centromeric regions in *A. thaliana* [18, 19], therefore we only aimed to assemble each chromosome arms in this study. In this context, our algorithm assembled each linkage group into a continuous sequence. In total, we were able to identify a telomer motif in eight assembled chromosome-arm sequences. Interestingly, only two telomer motifs have been observed in the Col-0 reference genome, indicating possible missing sequences in the reference genome.

In summary, our final assembly contains ten complete chromosome-arm sequences and three unanchored contigs. We also further analysed the three unanchored contigs and all of them were mapped to the centromeric region in in reference genome. Thus, the successful assembly of the *A. thaliana* genome by combining Pacbio and Nanopore sequencing data indicates that GALA provides a flexible framework for integrated assembly of multiple sequencing platforms.

### Assembly of a human genome

We next assembled a human genome using high-fidelity (HiFi) long reads generated by Pacbio using the circular consensus sequencing (CCS) mode [20]. For simplicity, we used the published assembly by canu [20] and the current human reference genome GRCh38.p13 as input of GALA. As the input reference genome is different from the genome to be assembled, GALA essentially created a reference-guided *de novo* assembly. The raw reads and the canu assembly were partitioned into 23 independent linkage groups. Each linkage group was then assembled with GALA. The comparison between our assembly and the published assembly can be found in **Fig.4a**. Overall, our assembly comprised of 37 continuous contigs, including 8 telomer-to-telomer gap-free pseudomolecular sequences, 4 near complete chromosomes each with a small telomeric fragment unanchored, 3 chromosomes with gapped centromeric regions. For 5 of the remaining chromosomes, we only assembled their long arms and the sequences for their short arms are too challenging for sequencing as they are also missing in the reference genome.

We illustrated our assembly chromosome-by-chromosome in **Fig.4b**. Two chromosomes worth mention here. In the reference genome GRCh38.p13 and also the published canu assembly, Chromosome 16 has several gaps and un-anchored contigs. GALA successfully assembled the chromosome into a single contig free of gaps of a length of 88Mbp. The assembled Chromosome 16 has a telomeric region at both ends, which is missing in GRCh38.p13. The second example is chromosome X, whose assembly is regarded as highly challenging and extra effort has been devoted to it in a recent paper. Our assembly with Chromosome X only contains two short gaps (about 0.75Kbp and 1.8Kbp) when compared to the recent published one [21]. The successful assembly of the human genome indicates that GALA can efficiently be applied to Pacbio HiFi data.

### Effect of the sequencing depth on the performance of GALA

We investigate how the performance of GALA changes with the data availability in terms of sequencing depth. This could be pivotal for experiment design considering the financial stress facing to most scientific research. We subsampled the original *C. elegans* Pacbio sequencing data to 20X, 30X, 40X, 50X, 60X, 70X, 80X, 90X, 100X, and 150X coverage, together with Hi-C data, and performed *de novo* assembly independently. Preliminary assemblies were generated using Canu, Flye, Mecat2, Miniasm and Wtdbg2 with raw and correctly reads. A detailed comparison can be found in Fig.5 and supplementary table 1. We have two interesting observations. At first, the gap-free *de novo* assembly is not a sensible option when the data coverage is less than 40X due to the limitation of current de novo assembly tools. As a consequence, GALA switches to gapped assembly for this scenario. Secondly, without Hi-C for scaffolding, Flye achieves the performance curve plateau at 60X coverage and GALA 40X coverage in terms of number of scaffolds and N50 of their assemblies. When Hi-C data is applied, the performance curve plateau starts from 40X for Flye and 20X for GALA (**Fig. 5a** and **b**) if very small gaps are allowed (<1Kbp). The higher coverage leads to better assembly for Flye with or without Hi-C data by lowering down the number of big gaps and misassemblies but has no notable effects on N50 and number of scaffolds (**Fig. 5c**). The higher coverage of data has no notable effect with GALA assembly in general.

**Figure 5.**
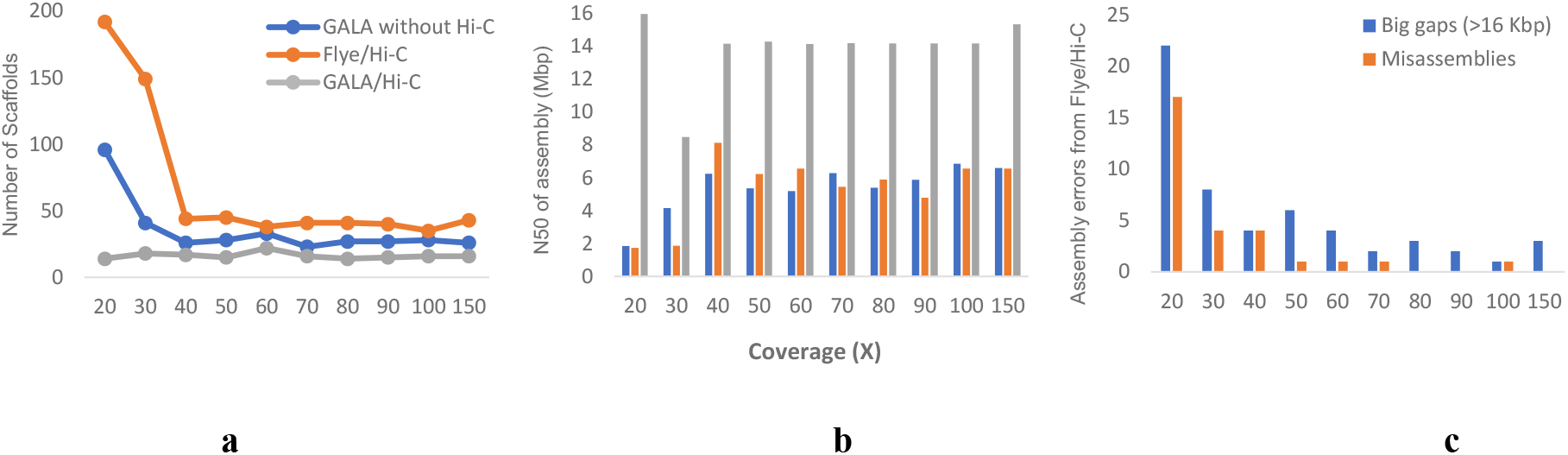
The assembly performances of GALA and Flye with Pacbio sequencing data at various coverages. Three assembly procedures have been presented: GALA without Hi-C data, Flye/Hi-C, GALA/Hi-C. The assembly performances are evaluated in terms of (a) the number of scaffolds, (b) N50, and (c) the number of big gaps (>16Kbp) and mis-assemblies. For simplicity, only the number of gaps and mis-assemblies for Flye/Hi-C have been shown, as only one misassembly has been identified in the assembly by GALA using 30X coverage sequencing data without the application of Hi-C data.

The performance of GALA, as well as almost all assembly software tools, changes significantly with raw reads length and sequencing error. Note that the above analyses are based on the Pacbio sequencing data generated with Pacbio RSII. As a consequence, the lengths of the raw reads are notably smaller and sequencing error is significantly higher than the current Pacbio Sequel II. In practice, the sequencing length distribution often vary significantly between different sequencing platforms, genome centres and sample preparation. It is difficult to set a straightforward threshold value for the minimum coverage of data for GALA assembly. As a thumb of rule, GALA can produce gap-free assembly from 25X coverage of Pacbio Sequel II data or Nanopore minION data if N50 of the raw data is larger than 20Kbp. For Pacbio HiFi, 20X coverage works well for GALA due to its low sequencing error rate.

### Effect of chromosome-by-chromosome assembly on the assembly graph

We investigated why GALA achieved complete assembly while existing tools had failed. We postulated that the chromosome-by-chromosome assembly strategy had played a role, and thus compared our assembly of *C. elegans* to this from Miniasm. We found a much simpler computer graph in the chromosome-by-chromosome assembly. In terms of the number of overlaps between reads (graph edges) in the assembly of *C. elegans*, the whole genome assembly generated 190,936,281 edges whereas the chromosome-by-chromosome assembly only generated 138,678,842 edges (27.37% less). A comparison between the chromosome-by chromosome assembly and the whole genome assembly is exemplified in **Fig. 6**.

**Figure 6.**
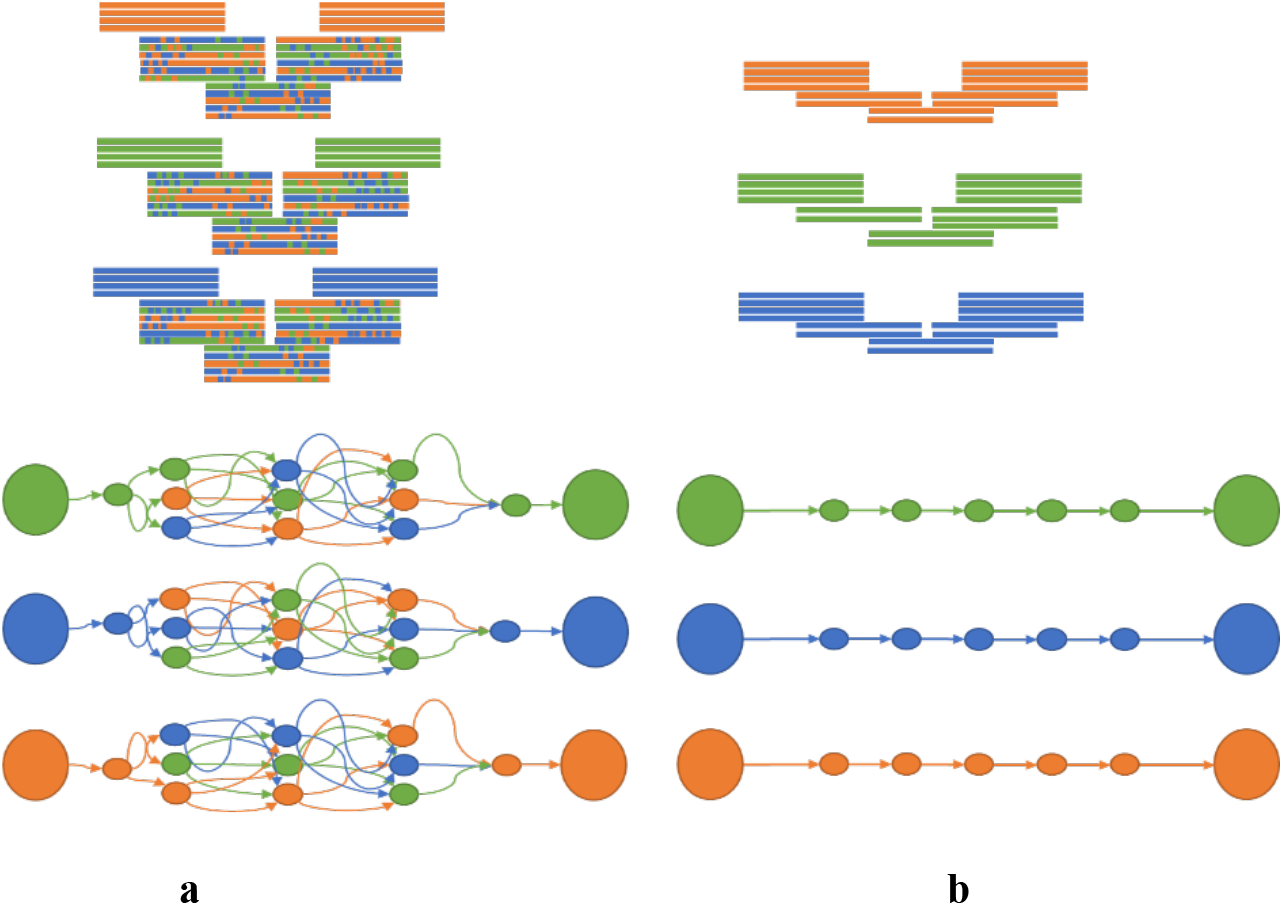
Comparison of the overlap graphs used by Miniasm during assembly of a region in the *C. elegans* genomes when the chromosome-by-chromosome strategy is applied or not. (a) In the whole-genome assembly mode, the overlap graph used by Miniasm contains numerous edges and extra effort is needed to collapse edges. (b) The chromosome-by-chromosome assembly allows a linear overlap graph to be derived by Miniasm in the same region.

The advantage of chromosome-by-chromosome assembly is more obvious in the regions which contain highly similar sequences, but still have unique markers, e.g. regions with ancient transposons (**Fig. 6)**. In addition, the regions which contain repetitive sequences but expanded by long reads usually allow a complete assembly by overlap graph-based algorithms, such as Canu or Mecat, but turn out to be too challenging to *de Bruijn* graph-based algorithms, such as Wtdbg2. In both scenarios, the GALA can obtain superior results.

## Discussion

In this paper, we have detailed GALA, a scalable method for gap-free chromosome-scale assembly. In many settings, GALA improved the contiguity and completeness of genome assembly compared to existing state-of-art assembly workflows and computational tools. Our new method is highly modular. In detail, the mis-assembly detection module (MDM) should be applicable for error correction regardless of the specific algorithm used for assembly; and the contig-clustering module (CCM) provides wide application in generating consensus assembly from multiple sequences. Although we have focused on *de novo* assembly, the modules in GALA should also work equally well in other applications.

GALA presents considerable advantages over the existing assembly workflows and computational tools. In this study, we generated chromosome-scale gap-free assemblies in most of our experiments. However, in certain circumstances, we failed to assemble a challenging region, such as centromeres in *A. thaliana* and also in the human genome. The failure is mainly due to the absence of raw sequencing data in that region (supplementary Fig. 5). Of note is that this failure also occurred in most of other commonly used computational tools too [19, 22–24]. The strength of GALA comes from the multi-layer computer graph model, which is highly flexible in incorporating heterogenous information. As clearly demonstrated in the assembly of the *C. elegans* and *A. thaliana* genomes, combinatory analyses of Pacbio and Nanopore sequencing data were achieved.

The performance of our GALA method also reflects the advantage of chromosome-by-chromosome assembly. Notably, the concept of chromosome-by-chromosome assembly was successfully tested on wheat genome assembly, for which expensive devices and time-consuming procedures have had to be applied [25, 26]. GALA is the first method to demonstrate that this can be achieved computationally. Finally, the concept of chromosome-by-chromosome assembly can also be applied to existing computational tools to refine an existing assembly.

## Methods

### Reciprocal alignment between preliminary assemblies

Minimap2 [27] (-x asm5) was used to map preliminary assemblies against each other. The raw and corrected reads were aligned to an assembly using BWA-MEM [28] with default parameters.

### Mis-assembly detection module (MDM)

We built a multi-layer graph by encoding the information from various preliminary assemblies *D*_*n*_. Each preliminary assembly *D*_*x*_ represented a layer that consists of a set of nodes *C*_*m*_, each node representing an assembled contig. The starting point of the MDM was reciprocal alignment of *D*_*n*_, which produced *n* * (*n* − 1) mapping files. We filtered the mapping files based on four criteria: (I) mapping quality (default 20), (II) contig length (default 5kbp), (III) alignment block size (default 5kbp), and (IV) sequence similarity percentage (default 70%). All parameters are tunable in GALA. A simple merging procedure was performed to merge nodes within the same layer if they are overwhelmingly supported by the reciprocal alignment to reduce the burden on computational resources.

We then linked the nodes between different layers by retrieving the information from reciprocal alignment. Assuming that a contig in node *C* in query layer *D*_*x*_, denoted as 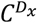, is mapped to a set of nodes in layer (*D*_*1,.n*_), denoted as 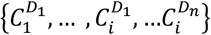, a mis-assembly at region *M* occurs if and only if contig 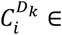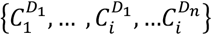 is partially mapped to 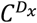.

Let *L* be the length of contig 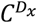, *N*_*A*_ be the number of contigs partially mapped to *M*, *N_B_* the number of contigs with complete alignment, and *N_S_* be the number of contigs starting or ending at *M*. We considered *M* as a genuine mis-assembled locus if:

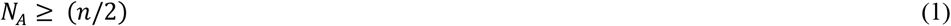

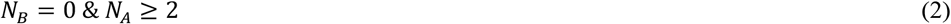

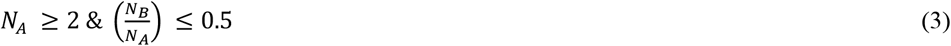

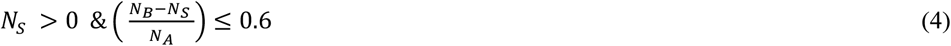

If a mis-assembly is identified, the node is split into two from the region *M*. This procedure iterates until the whole graph is free of mis-assemblies.

### Contigs clustering module (CCM)

The multi-layer computer graph output by MDM was expanded by adding into an extra layer representing the raw reads. So far, within each layer, nodes were separate from each other and no intra-layer edge existed. We first built intra-layer edges by browsing through the existing cross-layer edges. For node 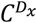 and its linked cross-layer node 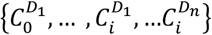, CCM starts by visiting all 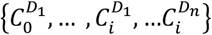. An intra-layer edge was built up if more than one node in the same layer was linked to the same cross-layer node. CCM pooled all connected nodes into a linkage group.

In the previous step of MDM, only contigs with a length larger than a certain threshold value, 5kbp at default, were encoded into our computer graph. Thus, those with smaller sizes were not used for mis-assembly detection. To avoid the situation where unique sequences could be missed out by accident, we kept them and classified them into existing linkage groups for further analysis.

If Hi-C information or a genetic map is available, extra links can be created between internal nodes. This approach would essentially lead to the merging of multiple independent linkage groups. CCM could also be performed in an iterative mode together with the linkage group assembly module (LGAM) as demonstrated in the examples below.

### Linkage group assembly module (LGAM)

The reads within a linkage group were assembled using assembly tools, e.g., Flye, Mecat and Miniasm. We mapped the newly assembled contigs against one of the chosen preliminary assemblies recorded in the same linkage group (free of mis-assembly) and removed those absent or those with multiple copies. A simplified version of the overlap graph-based merging algorithm was used to merge two contigs if necessary.

### *Caenorhabditis elegans* assembly

The Pacbio dataset contains three different runs and there was a clear batch effect with the sequencing quality and the amount of data between runs. We thus tested the assembly tools with either all runs (290X in coverage), or the biggest run alone (240X in coverage). We also used the reads-correcting-and-trimming module from Canu 1.8 [4] to correct the raw reads if the assembly tools take corrected reads as input. Preliminary assemblies were generated using Canu 1.8, Mecat2/Necat [3], Flye 2.4 [5], Miniasm 0.3-r179 [29] and Wtdbg2 [30], from Pacbio raw and corrected reads as well as nanopore raw reads. By comparing the summary statistics of preliminary assemblies, ten preliminary assemblies were chosen for GALA.

GALA modelled the preliminary assemblies and raw reads into 14 independent linkage groups. Seven of them were short continuous contigs and the others represented individual chromosomes or chromosome arms. Further analyses by blasting the seven short contigs in the NCBI database indicated that three of them were from *E. coli* contamination and one from the *C. elegans* mitochondrial genome, and thus were excluded from the subsequent analyses. The remaining three short contigs, two of them can be reliably put into the seven previously created linkage groups with the help of the assembly of Nanopore reads with Miniasm (**Supplementary Fig. 1**).

We assembled seven linkage groups with LGAM, each into a continuous sequence. Among the seven continuous sequences and one unanchored short contig, four of them revealed clear telomere repetitive motif at both terminals, indicating they are complete assemblies of single chromosomes. One chromosome-scale sequence had a telomere repetitive motif at one end, and its missing telomeric repetitive motif can be identified in the unanchored short contig, indicating they both should be merged as a single linkage group. The remaining two had a telomer repetitive motif at either side and their sizes clearly indicated they were two arms from a single chromosome. We thus pooled their linkage groups together. Finally, we re-assembled the two newly created linkage groups, and we were able to create complete sequences for the two chromosomes with a telomeric repetitive motif at both terminals. Further analyses indicated the split of this single chromosome into two linkage groups in the first run was mainly due to several tandem repeats.

### *Caenorhabditis elegans* genome assembly polishing and quality control

For a more accurate comparison, we polished our assembly with Pacbio and Illumina sequencing data. For this purpose, we first ran racon [31] with corrected Pacbio reads. The assembly was then polished using quiver 2.3.2 [32] with Pbmm2 1.1.0 as aligner. Finally, we ran pilon 1.23 [33] using Illumina sequencing data to correct short errors, especially those in homomorphic regions.

We evaluated the completeness of our polished assembly with Busco 3.0.0, and compared it to the published assembly, which is also polished using the same Illumina sequencing data, and the reference genome (**Supplementary Fig. 6**). We also aligned the Illumina short reads to our assembly using BWA-MEM and called the variants using BCFtools 1.9. Finally, we collected the statistics and compared them to those from the published assembly as a benchmark for the precision of the assembly.

### Arabidopsis *thaliana* assembly

Pacbio and Nanopore raw reads datasets were corrected using the Canu 1.8 correcting and trimming module. We found that Mecat2/Necat showed good performance for read correction in this dataset, so we additionally generated corrected Pacbio reads with Mecat2 with the minimal length of reads parameter as 5000 or 3000 separately, and corrected Nanopore reads with Necat with the minimal length of reads parameter 500. The preliminary assemblies were generate using Canu, Flye, Mecat2/Necat, Miniasm and Wtdbg2, from corrected and raw reads in both datasets. Thirteen preliminary assemblies were chosen for GALA with default parameters. In LGAM, we used Flye, Mecat2/Necat, Miniasm and Wtdbg2 tools.

## Declarations

### Ethics approval and consent to participate

Not applicable.

### Consent for publication

Not applicable.

### Availability of data and materials

#### Data Availability

Genome assemblies that were generated by GALA are available at https://doi.org/10.5281/zenodo.3840274. *C.elegans* Pacbio, Nanopro sequencing data and Illumina reads are available at PRJNA430756. *A.thaliana* KBS-Mac-74 Pacbio and Nanopro sequencing data are available at PRJEB21270. We downloaded Human rel3 Nanopore dataset from the Amazon Web Services Open Data https://github.com/nanopore-wgs-consortium/NA12878 and Humans HiFi dataset from https://obj.umiacs.umd.edu/marbl_publications/hicanu/index.html.

#### Code Availability

The source code of GALA is available from github at https://github.com/ganlab/gala. External software used in the current study were downloaded from the following URLs: Bcftools1.9, https://github.com/samtools/bcftools/releases; Busco 3.0.0, https://busco-archive.ezlab.org/v3; BWA 0.7.15-r1140, https://github.com/lh3/bwa; Canu 1.8, https://github.com/marbl/canu; Flye 2.4, https://github.com/fenderglass/Flye; Hifiasm 0.5-dirty-r247, https://github.com/chhylp123/hifiasm; IMR/Denom 0.5.0, http://chi.mpipz.mpg.de/imrdenom; MECAT2, https://github.com/xiaochuanle/MECAT2; Miniasm 0.3-r179, https://github.com/lh3/miniasm; Minimap 0.2-r124-dirty https://github.com/lh3/minimap; Minimap2 2.17-r941, https://github.com/lh3/minimap2; NECAT, https://github.com/xiaochuanle/NECAT; pbmm 2 1.1.0, https://github.com/PacificBiosciences/pbmm2; pilon 1.23, https://github.com/broadinstitute/pilon/releases; quiver 2.3.2, https://github.com/PacificBiosciences/GenomicConsensus; Racon 1.3.1, https://github.com/lbcb-sci/racon; SALSA2, https://github.com/marbl/SALSA; and Wtdbg2 2.5, https://github.com/ruanjue/wtdbg2.

## Authors’ contributions

XG conceived the project and interpreted the data. MA developed the GALA program and analyzed the data. XG and MA wrote the manuscript. The authors read and approved the final manuscript.

## Acknowledgements

We thank M. Tsiantis and R. Mott for their helpful comments on the work and Yuxia He for technical support. We also wish to acknowledge S. Morishita for sharing the original *C. elegans* Pacbio data with us.

## Reference

1. Cao, M.D., et al., Scaffolding and completing genome assemblies in real-time with nanopore sequencing. Nat Commun, 2017. 8: p. 14515.

2. Li, C., et al., Genome Sequencing and Assembly by Long Reads in Plants. Genes (Basel), 2017. 9(1).

3. Xiao, C.L., et al., MECAT: fast mapping, error correction, and de novo assembly for single-molecule sequencing reads. Nat Methods, 2017. 14(11): p. 1072–1074.

4. Koren, S., et al., Canu: scalable and accurate long-read assembly via adaptive k-mer weighting and repeat separation. Genome Res, 2017. 27(5): p. 722–736.

5. Kolmogorov, M., et al., Assembly of long, error-prone reads using repeat graphs. Nat Biotechnol, 2019. 37(5): p. 540–546.

6. Ellison, C.E. and W. Cao, Nanopore sequencing and Hi-C scaffolding provide insight into the evolutionary dynamics of transposable elements and piRNA production in wild strains of Drosophila melanogaster. Nucleic Acids Res, 2020. 48(1): p. 290–303.

7. Jiao, W.B., et al., Improving and correcting the contiguity of long-read genome assemblies of three plant species using optical mapping and chromosome conformation capture data. Genome Res, 2017. 27(5): p. 778–786.

8. Jibran, R., et al., Chromosome-scale scaffolding of the black raspberry (Rubus occidentalis L.) genome based on chromatin interaction data. Hortic Res, 2018. 5: p. 8.

9. Stankova, H., et al., BioNano genome mapping of individual chromosomes supports physical mapping and sequence assembly in complex plant genomes. Plant Biotechnol J, 2016. 14(7): p. 1523–31.

10. Song, B., R. Mott, and X. Gan, Recovery of novel association loci in Arabidopsis thaliana and Drosophila melanogaster through leveraging INDELs association and integrated burden test. PLoS Genet, 2018. 14(10): p. e1007699.

11. Chen, X. and M. Tompa, Comparative assessment of methods for aligning multiple genome sequences. Nat Biotechnol, 2010. 28(6): p. 567–72.

12. BSong B, S.Q., Wang H, Pei H, Gan X and Wang F, Complement Genome Annotation Lift Over Using a Weighted Sequence Alignment Strategy. Front. Genet, 2019. 10.

13. Bickhart, D.M. and G.E. Liu, The challenges and importance of structural variation detection in livestock. Front Genet, 2014. 5: p. 37.

14. Denton, J.F., et al., Extensive error in the number of genes inferred from draft genome assemblies. PLoS Comput Biol, 2014. 10(12): p. e1003998.

15. Zhang, X., J. Goodsell, and R.B. Norgren, Jr., Limitations of the rhesus macaque draft genome assembly and annotation. BMC Genomics, 2012. 13: p. 206.

16. Yoshimura, J., et al., Recompleting the Caenorhabditis elegans genome. Genome Res, 2019. 29(6): p. 1009–1022.

17. Ghurye, J., et al., Integrating Hi-C links with assembly graphs for chromosome-scale assembly. PLoS Comput Biol, 2019. 15(8): p. e1007273.

18. Michael, T.P., et al., High contiguity Arabidopsis thaliana genome assembly with a single nanopore flow cell. Nat Commun, 2018. 9(1): p. 541.

19. Pucker, B., et al., A chromosome-level sequence assembly reveals the structure of the Arabidopsis thaliana Nd-1 genome and its gene set. PLoS One, 2019. 14(5): p. e0216233.

20. Nurk, S., et al., HiCanu: accurate assembly of segmental duplications, satellites, and allelic variants from high-fidelity long reads. bioRxiv, 2020.

21. Miga, K.H., et al., Telomere-to-telomere assembly of a complete human X chromosome. Nature, 2020.

22. Arabidopsis Genome, I., Analysis of the genome sequence of the flowering plant Arabidopsis thaliana. Nature, 2000. 408(6814): p. 796–815.

23. Zapata, L., et al., Chromosome-level assembly of Arabidopsis thaliana Ler reveals the extent of translocation and inversion polymorphisms. Proc Natl Acad Sci U S A, 2016. 113(28): p. E4052–60.

24. Jiao, W.B. and K. Schneeberger, Chromosome-level assemblies of multiple Arabidopsis genomes reveal hotspots of rearrangements with altered evolutionary dynamics. Nat Commun, 2020. 11(1): p. 989.

25. Paux, E., et al., A physical map of the 1-gigabase bread wheat chromosome 3B. Science, 2008. 322(5898): p. 101–4.

26. Holusova, K., et al., Physical Map of the Short Arm of Bread Wheat Chromosome 3D. Plant Genome, 2017. 10(2).

27. Li, H., Minimap2: pairwise alignment for nucleotide sequences. Bioinformatics, 2018. 34(18): p. 3094–3100.

28. Li, H. and R. Durbin, Fast and accurate short read alignment with Burrows-Wheeler transform. Bioinformatics, 2009. 25(14): p. 1754–60.

29. Li, H., Minimap and miniasm: fast mapping and de novo assembly for noisy long sequences. Bioinformatics, 2016. 32(14): p. 2103–10.

30. Ruan, J. and H. Li, Fast and accurate long-read assembly with wtdbg2. 2019: p. 530972.

31. Vaser, R., et al., Fast and accurate de novo genome assembly from long uncorrected reads. Genome Res, 2017. 27(5): p. 737–746.

32. Chin, C.S., et al., Nonhybrid, finished microbial genome assemblies from long-read SMRT sequencing data. Nat Methods, 2013. 10(6): p. 563–9.

33. Walker, B.J., et al., Pilon: an integrated tool for comprehensive microbial variant detection and genome assembly improvement. PLoS One, 2014. 9(11): p. e112963.

