## Supplementary tables and figures for "GALA: gap-free chromosome-scale assembly with long reads" (from another version of this preprint: version 1, 2020-05-16)

**Supplementary Table 1.** The statistics *of C. elegans* preliminary assemblies**.** The column assembly indicates the names of assembly software tools used. Suffix (_Corr) indicates that the self-corrected sequencing data by canu has been used as input, otherwise, the raw data used. $^{+}$ 2 rounds of racon used to call the consensus from Miniasm;

| Platform | Assembly | Contigs | Genome Size | N50 | L50 | Longest Contig |
|---|---|---|---|---|---|---|
| Pacbio - all runs | **Canu** | 214 | 109,716,008 | 4,217,904 | 9 | 9,058,364 |
| | **Flye_Corr** | 68 | 104,652,135 | 3,693,393 | 9 | 12,185,728 |
| | **Miniasm_Corr$^{+}$** | 134 | 105,138,292 | 2,555,369 | 14 | 5,394,256 |
| | **Miniasm$^{+}$** | 469 | 115,467,549 | 2,387,630 | 17 | 4,834,144 |
| | **Wtdbg2** | 356 | 104,682,976 | 1,850,992 | 19 | 5,311,454 |
| | **Mecat_500** | 86 | 101,501,420 | 1,917,460 | 19 | 5,098,523 |
| Pacbio - one run | **Flye_Corr** | 41 | 102,134,972 | 4,201,097 | 9 | 10,503,810 |
| | **Wtdbg2_Corr** | 181 | 102,181,086 | 2,697,772 | 13 | 6,773,398 |
| Nanopore | **Flye** | 240 | 112,147,463 | 2,452,295 | 14 | 9,117,619 |
| | **Necat_5000** | 211 | 109,446,338 | 1,440,310 | 24 | 5,056,346 |
| | **Miniasm$^{+}$** | 574 | 138,450,772 | 640,109 | 59 | 2,924,146 |

**Supplementary Table 2.** The statistics of *A. thaliana* preliminary assemblies**.** The column assembly indicates the names of assembly software tools used. Suffix (_C_Corr) indicates that the self-corrected sequencing data by canu has been used as input, and suffix (_M_Corr) indicates that self-corrected sequencing data by Mecat as input, otherwise, the raw data used as input. $^{+}$2 rounds of racon used to call the consensus from Miniasm assemblies.

| Platform | Assembly | Contigs | Genome size | N50 | L50 | Longest Contig |
|---|---|---|---|---|---|---|
| Pacbio | **Canu** | 104 | 124,203,204 | 9,486,279 | 6 | 13,755,680 |
| | **Flye** | 115 | 120,306,161 | 14,783,068 | 4 | 16,179,764 |
| | **Flye_C_Corr** | 50 | 119,309,326 | 10,934,074 | 5 | 14,849,987 |
| | **Miniasm_C_Corr$^{+}$** | 67 | 120,798,896 | 9,142,171 | 6 | 13,769,655 |
| | **Wtdbg2_M2_Corr** | 165 | 120,851,855 | 11,409,806 | 5 | 13,011,670 |
| | **Mecat_5000** | 100 | 120,910,514 | 12,977,550 | 5 | 15,023,129 |
| | **Mecat_3000** | 70 | 120,856,793 | 11,625,809 | 5 | 13,685,097 |
| Nano | **Canu** | 288 | 113,408,765 | 6,522,919 | 7 | 12,090,110 |
| | **Fye_C_Corr** | 69 | 118,767,995 | 11,270,351 | 5 | 13,383,782 |
| | **Miniasm$^{+}$** | 67 | 118,382,021 | 11,760,723 | 5 | 14,259,632 |
| | **Miniasm_Minmap_C_Corr$^{+}$** | 55 | 116,067,126 | 5,914,303 | 8 | 11,151,688 |
| | **Wtdbg2_M2_Corr** | 220 | 119,593,115 | 8,796,241 | 6 | 13,156,465 |
| | **Necat_500** | 114 | 121,489,252 | 11,222,980 | 5 | 14,557,526 |

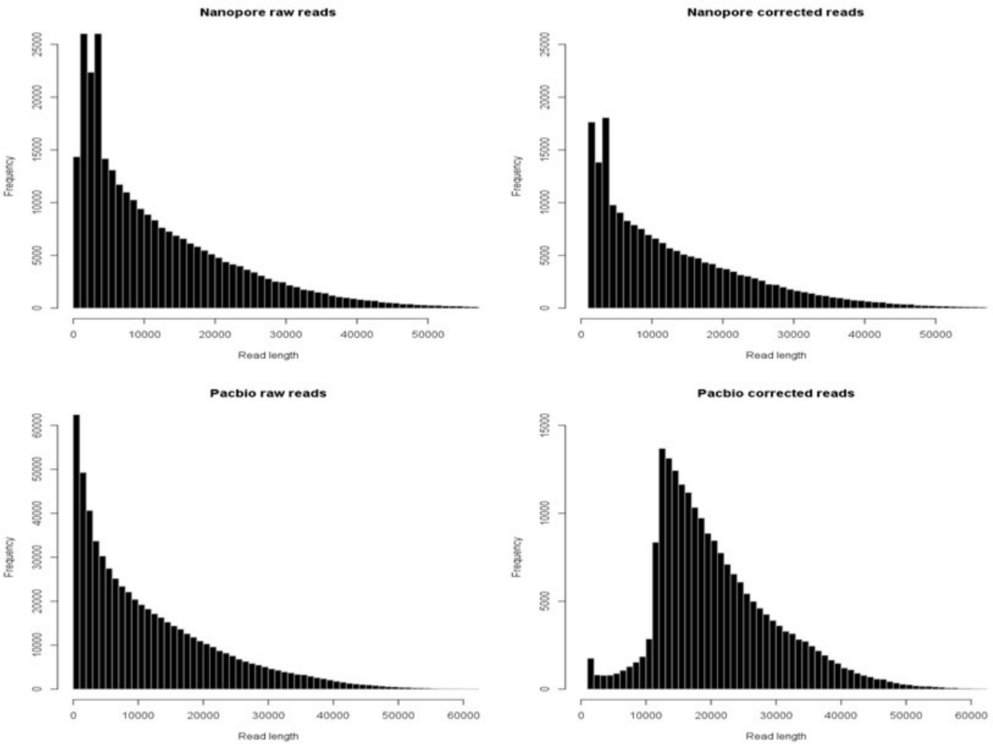


**Supplementary Figure 1.** The distributions of the length of the reads used for the assembly of *A. thaliana* genome. (a) Nanopore raw reads. (b) The self-corrected Nanopore reads by canu. (c) Pacbio raw reads. (d) the self-corrected Pacbio reads by canu.

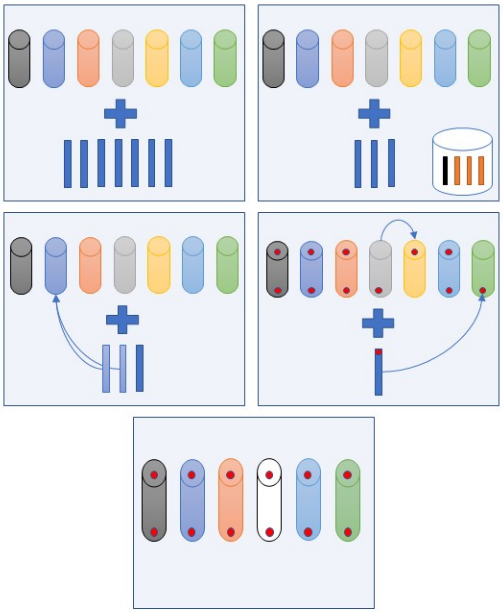

**Supplementary Figure 2.** The telomeric motif based analyses to merge several linkage groups in *C. elegans*. (a) GALA was applied on preliminary assemblies and raw reads, and produced seven scaffolding groups and seven short continuous contigs. (b) NCBI-blast showed that three contigs are from bacterial contamination (orange), a 13 kb mitochondrial genome (black). (c) Two of the remaining contigs were anchored to one of the linkage group by Miniasm/Nanopore assembly. (d) Four linkage groups had telomere motif at both terminals indicating the complete chromosome; two (grey and yellow) had only one telomere and were from a single chromosome indicated by their sizes; the last group (green) had only one telomere and the missing telomere appeared on the remaining short contig. (e) The chromosome-by-chromosome assembly successfully assembled each linkage group.

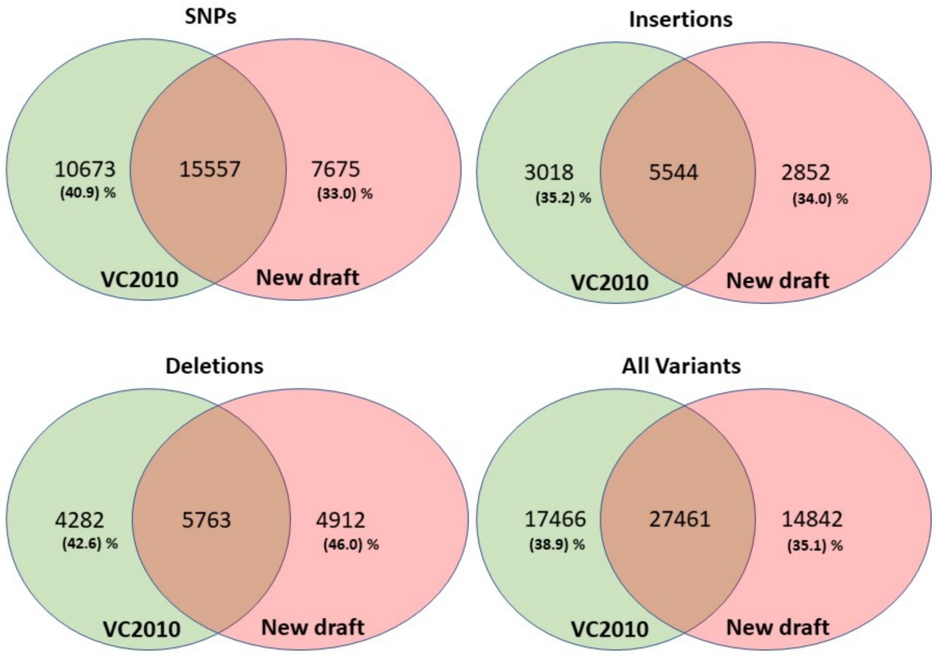


**Supplementary Figure 3.** Variant calling of *C.elegans* VC2010 and our new assembly against N2 reference. Bwa used for mapping and Denom [1] used for variant calling.

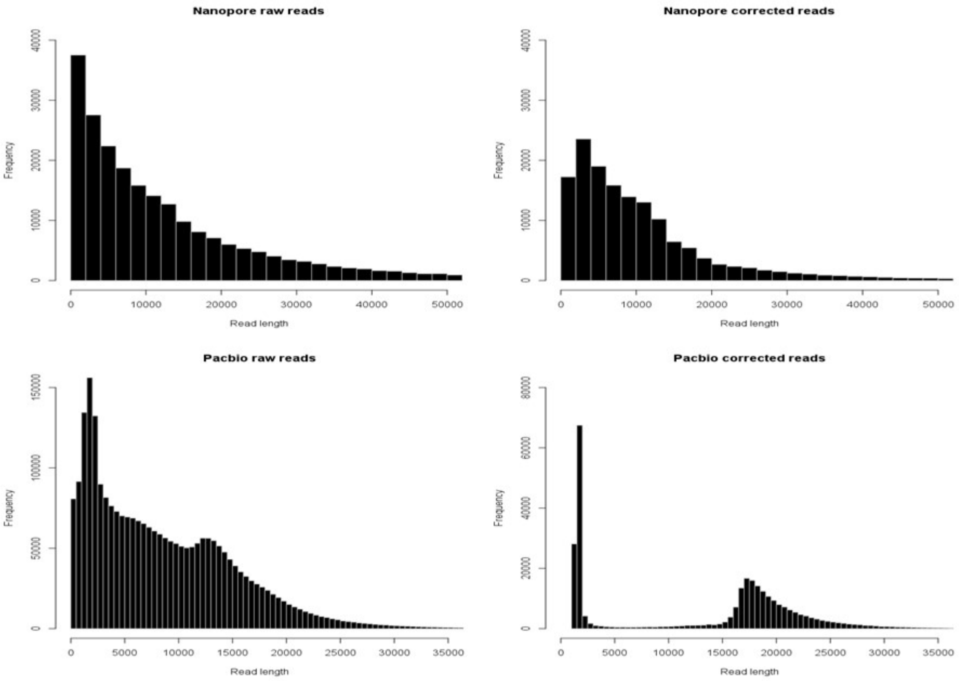


**Supplementary Figure 4.** The distributions of the length of the reads used for the assembly of *C. elegans* genome. (a) Nanopore raw reads. (b) The self-corrected Nanopore reads by canu. (c) Pacbio raw reads. (d) The self-corrected reads by canu.

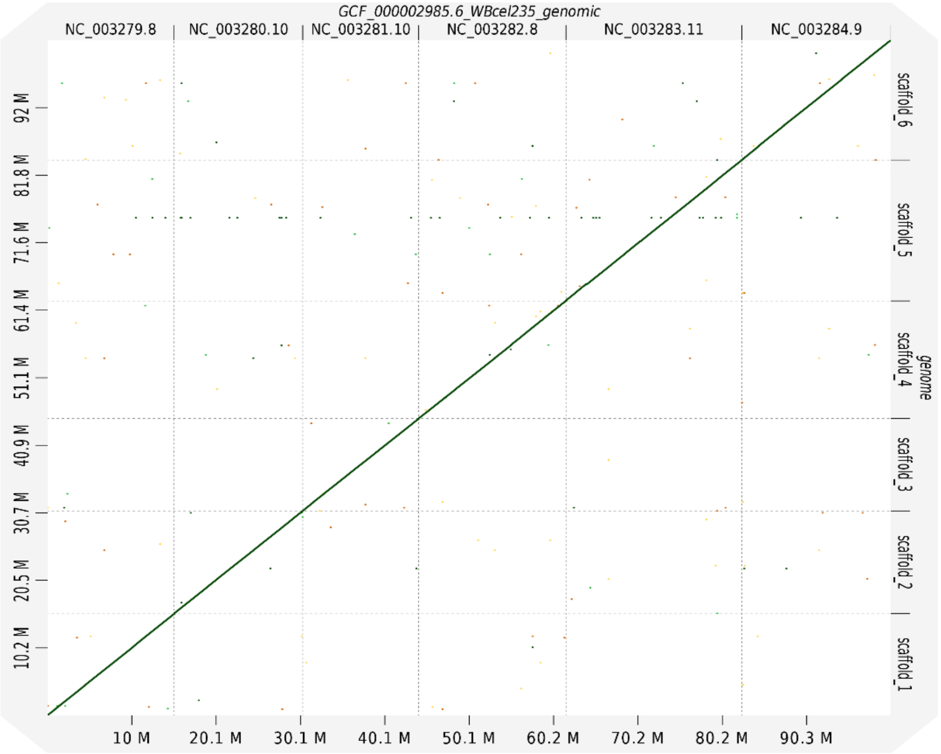


**Supplementary Figure 5.** Dotter plot of the alignments of our assembly to the *C. elegans* N2 reference genome.

**References:**


1. Gan, X., et al., *The Cardamine hirsuta genome offers insight into the evolution of morphological diversity.* Nat Plants, 2016. **2**(11): p. 16167.
